# Comprehensive genetic analysis by whole exome sequencing in 352 Korean pediatric patients with unknown neurodevelopmental disorders

**DOI:** 10.1101/472803

**Authors:** Youngha Lee, Jin Sook Lee, Soo Yeon Kim, Jaeso Cho, Yongjin Yoo, Sangmoon Lee, Taekyeong Yoo, Moses Lee, Jieun Seo, Jeongeun Lee, Jana Kneissl, Eunha Kim, Hyuna Kim, Woo Joong Kim, Jon Soo Kim, Jung Min Ko, Anna Cho, Byeong Chan Lim, Won Seop Kim, Murim Choi, Jong-Hee Chae

## Abstract

**Importance:** Accurate diagnosis of pediatric patients with complicated neurological problems demands a well-coordinated combination of robust genetic analytic capability and delicate clinical evaluation. It should be tested whether this challenge can be augmented by whole exome sequencing (WES).

**Objective:** To evaluate the utility of WES-based diagnosis and discovery of novel variants of undiagnosed patients with complex neurodevelopmental problems in a country with a centralized medical system.

**Design, setting, and participants:** A cohort of 352 Korean patients, believed to cover a major portion of the entire country from July 2014 to April 2017, with a broad spectrum of neurodevelopmental disorders without any pathogenic variants revealed by conventional methods were evaluated by trio-based WES at Seoul National University Children’s Hospital.

**Exposures:** WES of patients and parents and subsequent evaluation of genetic variants.

**Main outcomes and measures:** Genetic variants from each patient were evaluated for known disease association and novel variants were assessed for possible involvement with neurodevelopment process.

**Results:** We identified disease-causing variants, including newly discovered variants, in 57.4% of the probands, who had underwent a mean of 5.6 years of undiagnosed periods and visited mean of 2.3 tertiary hospitals. The cohort included 112 patients with variants that were previously reported as pathogenic (31.8%), 16 patients with copy number variants (4.5%) and 27 patients with variants that were associated with different clinical symptoms (7.7%). We also discovered potentially pathogenic variants from 47 patients that required further functional assessments (13.4%) and demonstrated potential implications in neurodevelopmental disorders. Following the genetic analysis, we provided more precise treatments to selected patients. A few clinical vignettes are presented that illuminate the potential diagnostic pitfalls that one could have encountered without this approach.

**Conclusions and relevance:** Our results highlight the utility of WES-based diagnosis for improved patient care in a country with a centralized medical system and discovery of novel pathophysiology mechanisms.

**Key points:** *Question:* What is the advantage of whole exome sequencing based diagnosis of pediatric neurology patients with unknown rare symptoms in a large tertiary clinic in a country with a centralized medical system?

*Findings:* Whole exome sequencing of 352 Korean patients, with a mean of 5.7 years of undiagnosed period, yielded 44.0% of conservative diagnostic yield. A number of cases were directly benefitted by trio-based WES via termination of diagnostic odyssey, genetic counseling for next offspring, or suggestion of more effective and customized treatment options.

*Meaning:* We report on the establishment of a national-level whole exome-based diagnosis system, with emphasis on deliberate integration of clinical interpretation and genetic analysis. Whole exome sequencing should be a choice of diagnostic tools for pediatric neurologic patients with ambiguous symptoms.

## Introduction

The inherent complexity of brain developmental processes inevitably leads to patients with diverse neurological problems that frequently challenge conventional diagnostic criteria. Despite recent improvements in methodologies for understanding genetic pathogenesis mechanisms, diagnosis of neurological disorders affecting children are frequently hampered by rarity and overlapping clinical features, which makes it difficult for clinicians to readily recognize the disease entity for proper treatment^1^. A genome-wide survey revealed that such gene-to-phenotype relationships still remain elusive for about half of human genes^2^. Due to the immense complexity of central nervous system development and the clinical observation that a large fraction of rare and undefined disease patients display neurological features^3,4^, it is reasonable to infer that pediatric neurologic patients are an impending target for the genome-wide genetic study. Indeed, large-scale systematic efforts have been conducted at regional or national scales^5–8^ to facilitate diagnosis and discovery of novel disease pathophysiology. As many rare neurologic disorders in children follow Mendelian inheritance, disease-causing variant discovery by trio-based whole-exome sequencing (WES) has been the most robust methodology, yielding an instant diagnosis rate of 25-41%^1,3,4,6,8^.

Importantly, the medical system in Korea provides a unique opportunity to conduct a systematic survey of rare disorders at a large scale. With a nationwide referral system toward a handful of major tertiary clinical institutions, Seoul National University Children’s Hospital (SNUCH) covers a large portion of the 51-million population, allowing for consistent evaluation and treatment of the patient cohort. For example, we recently reported genetic analyses of large patient cohorts of Duchenne muscular dystrophy (*n* = 507) and Rett-like syndrome without *MECP2* mutations (*n* = 34)^9,10^. Therefore, our study represents the largest of its kind conducted at a single clinic, with emphasis on the careful integration of clinical and genetic analyses.

Here we analyzed a cohort of 352 patients with various neurodevelopmental disorders using WES, experiencing a long undiagnosed period, and visiting multiple tertiary hospitals. We characterized the genotype-phenotype relationships of patients whose molecular defects were identified, and explored the potential association of genes with no previous disease associations. Notable case vignettes are introduced to further highlight the clinical utility of WES. Overall, we describe the establishment of a system that achieved an efficient integration of advanced genetic techniques in clinical diagnosis processes to maximize benefits for pediatric patients and their families in Korea.

## Methods

### Subjects

This study was approved by the Seoul National University Hospital Institutional Review Board (No. 1406-081-588). Blood samples were obtained from enrolled patients and their parents, who provided informed consent. WES was performed on 352 patients who visited the SNUCH pediatric neurology clinic from July 2014 to April 2017 and displayed various neurodevelopmental problems of unknown origins, such as demyelinating or hypomyelinating leukodystrophy, hereditary spastic paraplegia, mitochondrial disorders, epileptic encephalopathy, Rett syndrome-like encephalopathy, ataxia, neuropathy, myopathies, and multiple congenital anomalies/dysmorphic syndromes with developmental problems (Table 1). The patients can be categorized into two groups: (i) clinically diagnosable but with substantial genetic heterogeneity (82/352, 23.3%) or (ii) heterogeneous and nonspecific clinical presentations without definite diagnosis (270/352, 76.7%; eFigure 1 in the Supplement). Prior to the WES analysis, thorough clinical and laboratory evaluations and extensive patient examinations have been conducted to identify possible genetic causes. These included genetic tests with candidate gene sequencing, targeted gene sequencing panel, trinucleotide repeat analysis, microarray, metabolic work-up, brain/spine MRI, or muscle biopsy if applicable. All subjects were evaluated by three pediatric neurologists, two pediatric neuroradiologists, and a pathologist.

**Table 1.** Clinical information of 352 patients.

|  |  |
| --- | --- |
| Sex (*n* (%)) |  |
| Male | 159 (45.2) |
| Female | 193 (54.8) |
| Age at symptom onset (years) | 1.4 (0-21) |
| Age at first access to a tertiary hospital (years) | 1.8 (0-21) |
| Interval between symptom onset and first medical access (months) | 4 (0-136) |
| Number of visited tertiary hospitals for diagnosis (*n* (%)) |  |
| 1 | 38 (10.8) |
| 2 | 180 (51.1) |
| 3 | 114 (32.4) |
| 4 | 18 (5.1) |
| 5 | 2 (0.6) |
| Age at WES (years) | 7.5 (0-53) |
| Interval between the first access and WES (months) |  |
| Patients aged 0-10 years | 33.3 (1-92) |
| Patients aged >10 years | 114.6 (7-624) |
| Primary clinical diagnosis (*n* (%)) |  |
| Neurodegenerative disease | 66 (18.8) |
| Neurometabolic disorder | 46 (13.1) |
| Epileptic and developmental encephalopathy | 36 (10.2) |
| Hypotonia | 25 (7.1) |
| White-matter abnormalities | 22 (6.3) |
| Unknown spasticity | 15 (4.3) |
| Other | 142 (40.3) |
| Number of involved specialists for diagnosis (*n* (%)) |  |
| 1 | 116 (33.0) |
| 2-3 | 170 (48.3) |
| > 3 | 66 (18.8) |
| Straight-line distance from home to the clinic, km (*n* (%)) |  |
| < 20 | 118 (33.5) |
| 20-100 | 112 (31.8) |
| > 100 | 122 (34.7) |

### Whole Exome Sequencing

WES was performed at Theragen Etex Bio Institute (Suwon, Korea) following the standard protocol and the data were analyzed as described previously^10^. Depending on the genetic analysis result, each patient was categorized as one of the following: category 1: known disease-causing genes were found; category 2: causative gene for other diseases were found; category 3: potentially pathogenic gene, but without prior disease association, was found; category 4: no disease-causing candidates were found; and category 5: known pathogenic copy number variation (CNV) was found.

Our variant assessment procedures were as follows: firstly, patient-specific CNVs were checked and samples with CNVs were classified as category 5. Then, patient-specific nucleotide variants such as *de novo*, compound heterozygous, and rare homozygous variants were selected by comparing against parents and prioritized based on the inheritance pattern (Figure 1A). In correlating the patient’s symptoms with genetic variants, if patients carried a known pathogenic variant in OMIM or ClinVar, they were categorized as category 1 or 2, depending on similarity with reported and observed symptoms. For previously unreported variants, if they were never seen in normal individuals (Exome Aggregation Consortium (ExAC)^11^, 1000 Genomes Project^12^, Korean Variant Archive (KOVA)^13^ and in-house variants) and were evolutionarily well-conserved at the amino acid level, they were categorized as potentially pathogenic. For CNV calling, the normalized coverage depths of each captured intervals were compared to the depths from related individuals.

**Figure 1.**
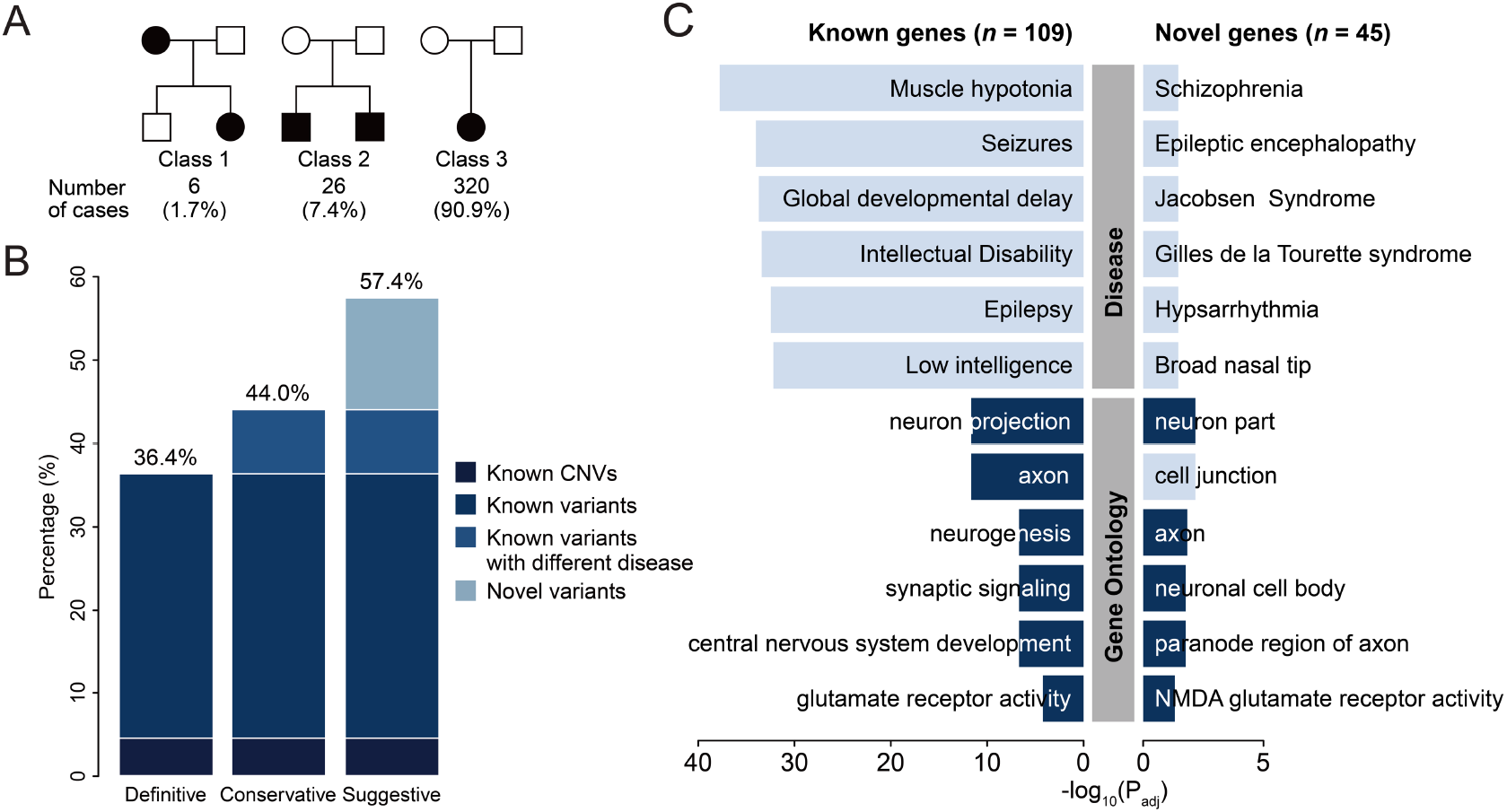
Classification of neurologic disease cohort and results of clinical WES analysis. **(A)** Subjects by disease inheritance patterns. Class 1: autosomal dominant families; Class 2: families with affected siblings; Class 3: affected individuals with no family history. **(B)** Diagnostic yields of 352 patients with undiagnosed symptoms using WES. **(C)** GO enrichment analysis of 109 known and 45 novel genes. Categories with dark blue colors denote GOs related to the nervous system.

### Human Brain Transcriptome Data

The BrainSpan transcriptome database (http://www.brainspan.org) has been used to build developing human brain networks^14^. BrainSpan data from eight post-conceptual weeks to 40 years of age were analyzed. A total of 385 samples were used for the analysis after combining the multiplicates by taking mean values. Probes with TPM (transcripts per kilobase million) > 5 in at least one sample were used, yielding 23,943 probes as “brain-expressed transcripts”.

### Brain Transcriptome Network Analysis

Using the above brain-expressed transcripts, we created eight known gene coexpression networks by selecting genes that are highly correlated to our 109 known genes (Pearson’s correlation coefficient > 0.7) from each developmental period (eFigure 2A in the Supplement). Then, we asked whether our novel genes can be successfully integrated into the known gene coexpression network. We randomly selected 45 genes (equal to the number of novel genes) in brain-expressed transcripts and counted how many edges they formed with the known genes. The 10^5^ random gene selections were performed and the number of edges with a known gene was used to construct a distribution. The number of edges from our observed novel genes was evaluated against the random distributions. P-values were calculated using z-score, assuming normal distrubutions.

## Results

### WES analyses yielded a diagnostic success rate of 44.0%

The symptoms of the patients were mostly of pediatric onset (mean 1.4 years of age), harbored neurodevelopmental problems, and patients were soon referred to tertiary hospitals (mean 1.8 years of age). The majority of the patients visited multiple tertiary hospitals for diagnosis (89.2% visited more than one hospital, mean of 2.3 hospitals), required a mean of 2.4 specialists (18.8% required more than three) and spent a mean of 5.7 years until being analyzed by WES at SNUCH (Table 1, eFigure 3 in the Supplement).

The majority of the patients occurred sporadically (320/352, 90.9%; Figure 1A), making them suitable for trio-based WES analysis. After integrative assessments of genetic variants, their clinical impacts, and patient symptoms, we were able to diagnose 36.4% (128/352) of the cohort with high confidence. The patients included carriers of CNVs (16/352 = 4.5%; 12 deletions and 4 duplications; eTable 1 in the Supplement), where 14 CNVs were originated *de novo* (4.0%), which is slightly lower but comparable to the previous observation^15^. Two inherited pathogenic CNVs were identified; a 165.5 kb deletion was discovered from a large family with multiple affected individuals (eFigure 4 in the Supplement) and a 299.5 kb hemizygous duplication was transmitted from a healthy mom to her affected son (eTable 1 in the Supplement). An additional 7.7% of the cohort (27 patients) harbored variants that were previously reported and assumed to be pathogenic, but displayed distinct phenotypes, potentially expanding the phenotypic spectrum of these genes. For example, two patients initially diagnosed with muscle hypotonia were found to carry a pathogenic heterozygous nonsense or missense variant in *COL1A1*, known to cause osteogenesis imperfecta^16^. They did not display skeletal problems, but showed blue sclera^17^. Adding this group yielded an instant diagnostic rate of 44.0% (“known genes”; eTable 2 in the Supplement).

Finally, 13.4% of the cohort (47 patients, 45 genes) harbored variants that are highly likely to be pathogenic but their disease associations are elusive (“novel genes”), yielding a “suggestive” diagnostic yield of 57.4% (Figure 1B). Overall, 58.6% of the variants are of *de novo* origin, and 22.9% are loss-of-function (LoF) to the gene (eFigures 5, 6 in the Supplement).

### Novel genes display potential enrichment in neuronal differentiation

We assessed if our 45 novel genes indeed posed a neurologic disease-causing function. Firstly, the novel genes displayed significant enrichment in gene ontologies (GOs) related to neurodevelopment, mimicking the pattern of known genes (Figure 1C; GO:0097458, neuron part, *P_adj_* = 6.7 × 10^−3^; GO:0030424, axon, *P_adj_* = 1.5 × 10^−2^). Since our cohort is composed of patients with diverse neurodevelopmental features, we split the cohort into several broad categories by presence or absence of epilepsy, severe developmental delay (DD) or facial dysmorphisms (FD). By evaluating gene groups that are enriched in each patient group, we identified genes that are involved in each clinical feature. A comparison of genes found from patients with or without epilepsy leads to the enrichment of synapse-related genes in epileptic patients (GO:0044456, synapse part, *P_adj_* = 4.2 × 10^−5^; GO:0099537, trans-synaptic signaling, *P_adj_* = 1.6 × 10^−4^), whereas patients without epilepsy showed myelination-related gene enrichment (Figure 2A; GO:0043209. Myelin sheath, *P_adj_* = 1.5 × 10^−6^). Patients with severe DD carried genes related to neurotransmitter signaling and hypotonia (Figure 2B; disease:C0026827, muscle hypotonia, *P_adj_* = 3.4 × 10^−24^; GO:0035235, ionotropic glutamate receptor signaling pathway, *P_adj_* = 1.3 × 10^−8^), and patients with FD displayed enriched genes in organ morphogenesis genes (Figure 2C; disease:C0424688, small head, *P_adj_* = 3.3 × 10^−7^; GO:0035295, tube development, *P_adj_* = 1.8 × 10^−2^). Applying this patient subgrouping strategy, we monitored if the diagnostic ratio would differ by presence or absence of such features. Remarkably, patients with FD showed a 17.9 % higher diagnostic ratio than those that do not, and other categories displayed similar patterns (Figure 2D-E).

**Figure 2.**
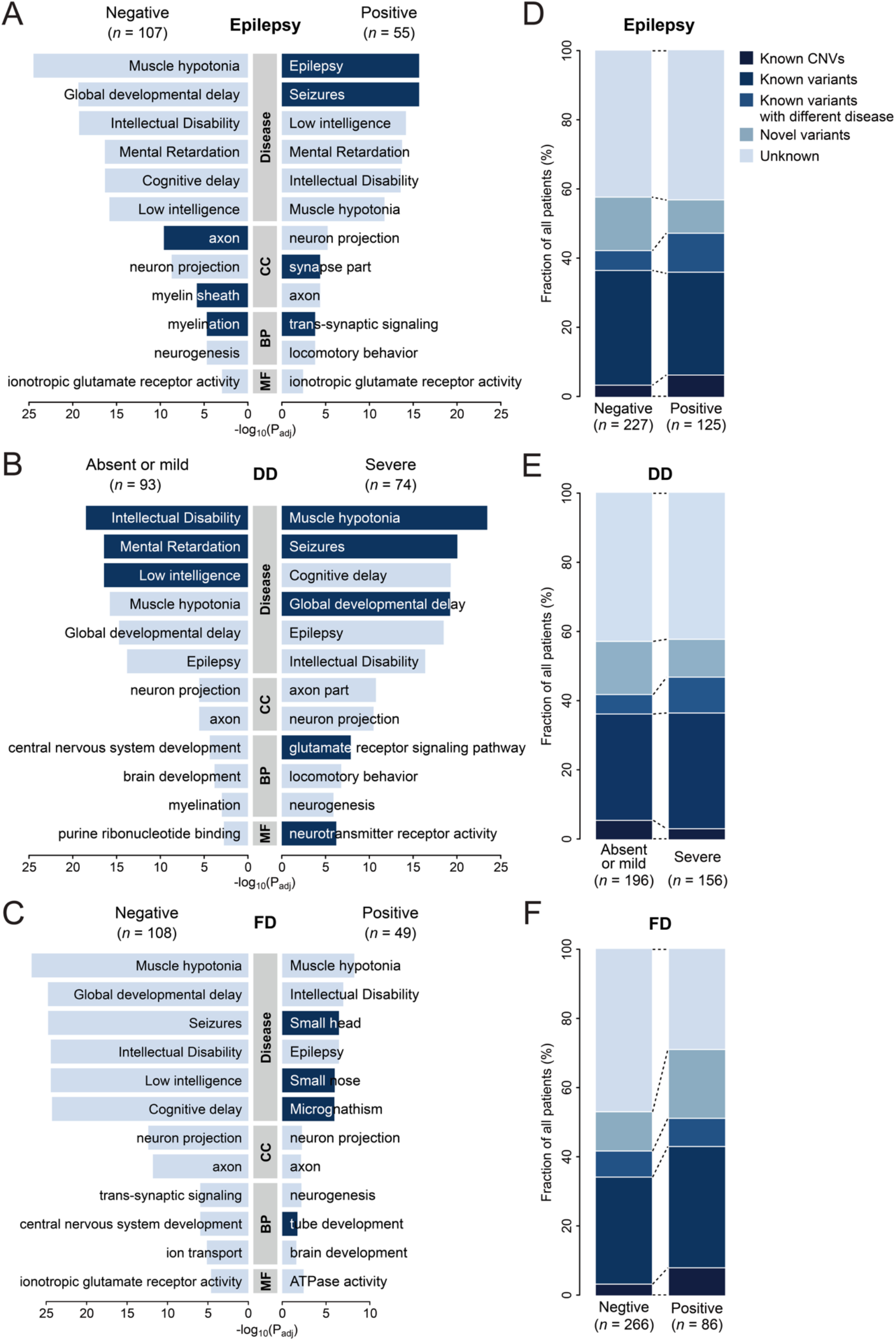
Patient subgroup analysis by major symptoms. (A-C) GO analysis of genes with pathogenic variants that were carried by patients with or without the epilepsy feature (A), with or without severe DD (B), and with or without notable FD (C). Categories indicated by a dark blue color denote the GO or disease features that are specifically enriched in each group. (D-F) Diagnostic yields according to the epileptic symptom (D), the severity of DD (E), and the notable FD (F). Disease, DisGeNET curated annotation; CC, Cellular Component of GO; BP, Biological Process; MF, Molecular Function; DD, Developmental delay; FD, Facial dysmorphism.

Next, we evaluated if our novel genes as a group displayed a strong correlation in expression with known disease-associated genes during brain development by simulating our novel gene set against the Allen Brain Atlas data (Materials and Methods). After 10^5^ permutations, we found that the observed novel gene involvement was significantly stronger compared to the randomly selected gene sets across eight developmental windows (eFigure 2B in the Supplement; all periods show *P_adj_* < 0.05). Furthermore, this test was expanded to the four anatomical domains in each period, yielding 32 spatio-temporal windows (eFigure 2C in the Supplement). It is notable that the most highly enriched windows are concentrated in the frontal cortex area (eFigure 2D in the Supplement; R1 × P1-4). The results suggest that our novel genes are expressed in concordance with known disease-causing genes in developing brain tissues and this phenomenon is concentrated in the frontal cortex.

### Notable vignettes

WES-based diagnoses of 352 patients with neuro-developmental problems generated cases that were clinically meaningful for a number of reasons (Figure 3, Table 2).

**Figure 3.**
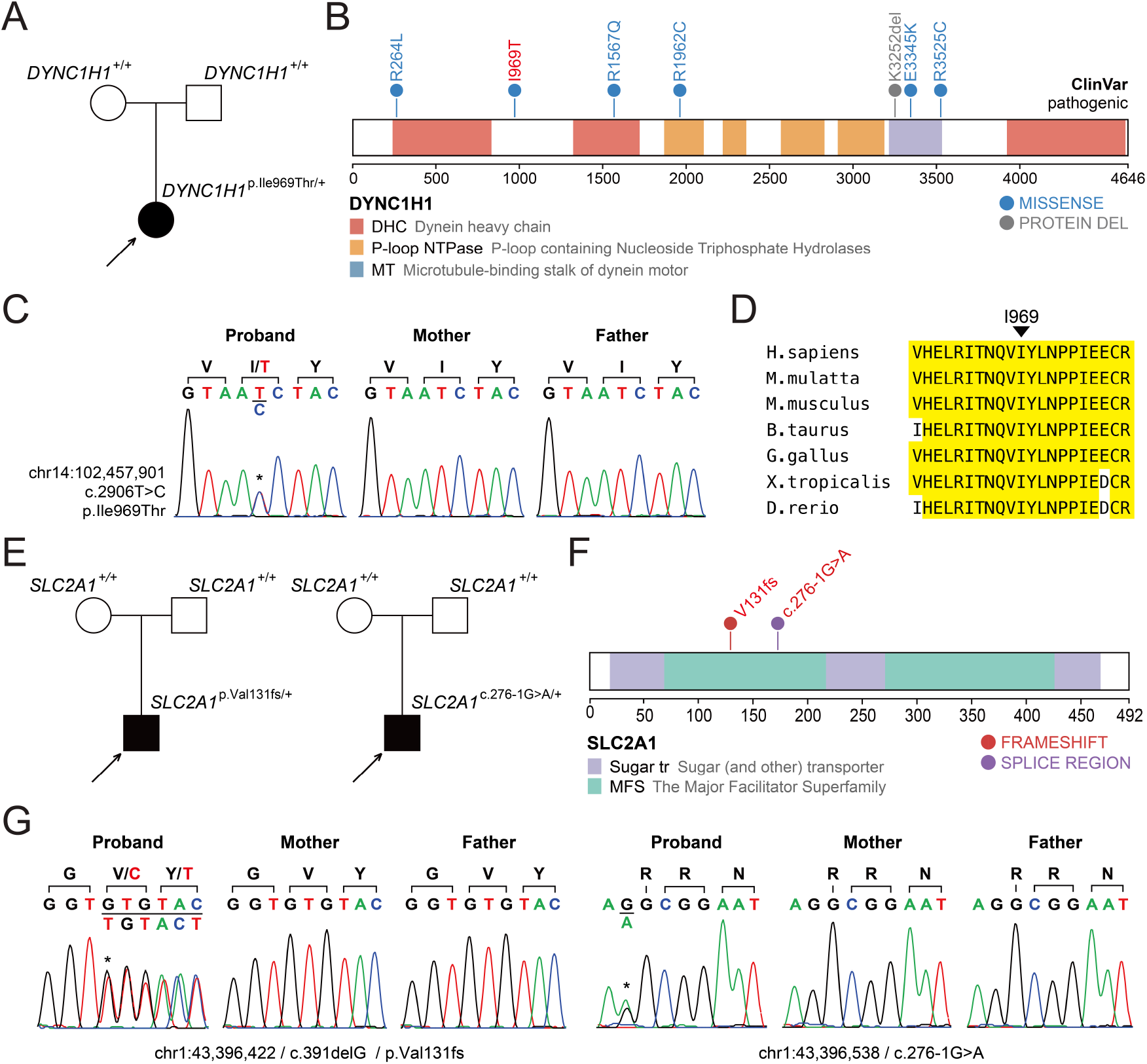
Presentation and validation of cases that WES-based analysis altered clinical courses. (A) A *de novo* variant in *DYNC1H1* identified from a patient with delayed development and hypotonia. (B) Pathogenic variants from ClinVar and domains of DYNC1H1 are displayed. A variant discovered in this study is shown in red. (C) Sanger traces validating the p.Ile969Thr variant. (D) Evolutionary conservation of the Ile969 residue across orthologs from major vertebrate species. (E) Two patients with dystonia and delayed motor development harbored LoF *de novo* variants in *SLC2A1*. (F) The variants found in two patients and domains of SLC2A1 are displayed. (G) Sanger traces validating the *de novo* variant.

**Table 2.** Notable cases where WES-based analysis conferred correct diagnoses or changed medical treatment strategies.

| Initial clinical problem | Causal gene | Modified clinical interpretation (MIM number) | Significance of WES-based patient evaluation (treatment) | References |
| --- | --- | --- | --- | --- |
| Developmental regression with RTT like phenotype | *ST3GAL5* | Salt and pepper developmental regression syndrome (#609056) | Identified the molecular defect and established an accurate diagnosis | 18,19 |
| Hypotonia and motor delay followed by lower extremity weakness | *DYNC1H1* | Spinal muscular atrophy, lower extremity-predominant 1, AD (#158600) | Diagnosed a case with pleiotropic and evolving symptoms | 20 |
| Early onset hypotonia, sacral mass, congenital heart disease, and facial dysmorphism | *ASAH1* | Farber lipogranulomatosis (#228000) | Corrected a misdiagnosis | 21 |
| Ataxia followed by generalized dystonia | *ANO3* | Expanded spectrum of dystonia 24 (#615034) | Suggested a treatment strategy that resulted in gradual improvement within one year (deep brain stimulation) | 30 |
| Focal lower leg dystonia, dystonic gait | *SLC2A1* | GLUT1 deficiency syndrome 2 (#612126) | Identified disease-specific treatment that resulted in near-elimination of dystonia (ketogenic diet) | 22 |
| Leigh syndrome | *SLC19A3* | Thiamine metabolism dysfunction syndrome 2 (#606152) | Identified disease-specific treatment that resulted in clinical improvements in dystonia, spasticity, and cognitive function (supplements of thiamine and biotin) | 31 |
| Recurrent infections, telangiectatic skin mottling, and brain infarctions | *TMEM173* | STING-associated vasculopathy, infantile-onset (#615934) | Provided a rationale for a new treatment strategy that improved the skin lesions (tofacitinib treatment) | 29 |
| Severe global developmental delay, seizures, and acanthotic skin lesions | *RAB11B* | Neurodevelopmental disorder with ataxic gait, absent speech, and decreased cortical white matter (#617807) | Identified a new disease gene leading to a neurodevelopmental syndrome | 23 |

### ST3GAL5 case: genetic elucidation of a case of compound symptoms^18^

Two female siblings presented with a Rett syndrome-like phenotype, such as psychomotor retardation/regression, delayed speech, hand stereotypies with a loss of purposeful hand movements, and choreoathetosis. Genetic tests of *MECP2/CDKL5/FOXG1* and chromosomal microarray found no plausible candidates. GM3 synthase deficiency with *ST3GAL5* compound heterozygous variants was diagnosed through WES, which was confirmed by liquid chromatography-mass spectrometry analysis^18,19^. WES ended three years of an undiagnosed period and broadened the phenotypic spectrum of this rare neurometabolic disorder.

### DYNC1H1 case: a case of evolving symptoms

An 8-month-old girl from healthy parents presented with global developmental delay and generalized hypotonia. Laboratory findings, MRIs, electromyography/nerve conduction studies, and genetic tests including *SMN1*, failed to identify her etiology. Although her cognitive function regression improved rapidly to near normal, a suspicious paraplegic gait developed from three years of age. WES revealed a novel *de novo* missense variant in *DYNC1H1*, encoding a subunit of the cytoplasmic dynein complex (Figure 3A-D). This observation led us to conclude that the patient started displaying global DD, but was eventually diagnosed with SMALED (spinal muscular atrophy, lower extremity-predominant), mimicking hereditary spastic paraplegia. This case poses a rare example, along with recent recognition that *DYNC1H1* carriers display complex HSP^20^, which demonstrates the clinical utility for pediatric cases with phenotypic pleiotropy and symptom evolution.

### ASAH1 case: a case with corrected diagnosis^21^

An 11-month-old girl from healthy parents was referred to our hospital for developmental delay with hypotonia, facial dysmorphism, congenital heart defects, and a sacral mass since birth. Karyotyping, metabolic screening, and chromosomal microarray revealed no abnormalities. Excision of the sacral mass led to pathologic diagnosis of epithelioid hemangioendothelioma, which required repeated operations and chemotherapy. However, joint contracture and multiple subcutaneous nodules appeared from 18 months of age. WES revealed compound heterozygous variants in *ASAH1*, associated with Farber disease. After the establishment of the correct molecular diagnosis at the age of four, additional electron microscopic findings of previously excised masses confirmed the pathognomonic findings^21^.

### SLC2A1 case: a case where an effective treatment was given

A 16-year-old boy from an otherwise a healthy family displayed an abnormal gait from around eight years of age that later developed into falls with dystonia, notably in the lower extremities, which was provoked mainly by exercise or stress (Video 1). His motor development was delayed, although he had normal head circumference and cognition. WES revealed a *de novo* LoF variant in *SLC2A1*, encoding a glucose transporter (GLUT1)^22^ (Figure 3E-G). We also retrospectively noticed that he had developed episodes of staring spells from an age of four years, suggesting absence seizures, which disappeared after three years of antiepileptic medication. Cerebrospinal fluid analysis confirmed a low glucose level (37 mg/dL, 36% of serum glucose, normal range > 40%). Based on this observation, a ketogenic diet was applied and completely changed the quality of his life, with a resulting near-absence of dystonia (Video 2).

### RAB11B case: a case of newly identified disease through international data sharing^23^

A 2-year-old boy was referred for evaluation of global developmental delay. He showed microcephaly and severe cognitive impairment with epilepsy but without evidence of regression. Notably, abnormal acanthotic skin lesions progressed from the face to the whole body. We identified a *de novo* variant in *RAB11B* and submitted this to GeneMatcher^24^. Soon, four cases with a similar neurodevelopmental phenotype were matched and we demonstrated that this gene causes intellectual disability and a distinctive brain phenotype^23^.

## Discussion

Our study demonstrated the clinical utility of WES in children with various and complex neurodevelopmental disorders. We identified genetic causes and sometimes corrected misdiagnoses to end the long journeys to diagnoses. We also found medically actionable cases where it was possible to offer the appropriate treatment options and a drug repositioning strategy (Table 2). As such, WES facilitates enhanced clinical care, genetic counseling and a reduction of the psychosocial stress and medical costs for patients and their families.

Consistent with the previous studies, we were able to diagnose about one third of the patients (Figure 1). Our approach allowed for an expanding phenotypic spectrum of known genes (27/352 cases, 7.7%), and suggested novel genes that may allow for a further understanding of neurodevelopmental disease mechanisms (47/352 cases, 13.4%). The novel genes showed significant enrichment in GOs related to neurodevelopment processes, and were concentrated in frontal cortex regions (Figures 1D and eFigure 5 in the Supplement). Furthermore, we observed differential diagnostic ratios when the cohort was subdivided into several major symptoms (Figure 2). Data sharing programs such as GeneMatcher have proven to be pivotal when assessing these novel variants^10,24^. The matching program allowed us to collect more patients of similar genotype and phenotype, and also provided evidence revealing whether the candidates are more likely to be true or not (eTable 3 in the Supplement).

Still, 42.6% of the cases (150/352) remained undiagnosed after our WES effort, suggesting a substantial opportunity for further improvement (Figure 1). Along this line, a systematic re-analysis effort with additional bioinformatics pipelines increased the diagnostic rate by 4.2%^25^. Also, searching for functional non-coding variants through whole genome sequencing (WGS) and evaluating multiple rare functional variants that may increase disease predisposition might be beneficial^26^, although a recent meta-analysis study claims only a minimal improvement in the diagnostic yield by WGS presumably due to our limited understanding of noncoding variant functions^27^. An alternative approach would be to cross-link genome data with transcriptome data, to identify functional cryptic variants that directly influence expression of critical genes^28^, although preparing patient-derived tissues still remains a practical challenge.

Nevertheless, we highlight the clinical challenges that were posed by an evolving phenotype over time in growing children and how this can be overcome (Figure 3A-D), which facilitates identification of treatable or actionable cases (Figure 3E-G). Our patient cohort also included a successful drug repositioning case for a rare neurogenetic disease^29^ (Table 2). All of these cases are expected to increase as more genotype-phenotype relationships are discovered and more drugs become available. The unique Korean medical system leads to the concentration of clinically challenging patients into a few tertiary clinics, although patients still frequently experience unnecessary referrals and repeated tests. This study demonstrates that applying WES and subsequent in-depth analysis provides clinical and practical benefits to patients and their families.

Our study presented a genetic analysis of 352 Korean pediatric patients with unexplained neurodevelopmental problems, revealing various known and novel genetic etiologies. We provided rationales to aggressively extend our system to a wider ranges of undiagnosed rare disease patients in countries with centralized healthcare like Korea. Meanwhile, we also improved genome data generation and analytic methods. Through a careful integration of detailed phenotyping, genetic analysis and data sharing, we demonstrate how this approach allowed for more precise diagnosis and personalized patient care. Finally, we reported on the successful establishment of such processes in Korea for patients with various undiagnosed neurodevelopmental disorders.

## Author contributions

Drs Chae and Choi had full access to the data in the study and take responsibility for the integrity of the data and the accuracy of the data analysis.

Study concept and design: Chae, Choi

Analyzing genome data: Y. Lee, Y. Yoo, S. Lee, T. Yoo, M. Lee, Seo, J. Cho, J. Lee, Kneissl, E. Kim, Choi

Providing clinical data: J. S. Lee, S. Y. Kim, H. Kim, W. Kim, J. Kim, Ko, A. Cho, Lim, W. S. Kim, J. Cho, Chae

Combining genetic and clinical data: J. S. Lee, Y. Lee, S. Y. Kim, Choi, Chae

Genetic and statistical evaluation of the cohort: Y. Lee, Choi

Drafting the manuscript: J. S. Lee, Y. Lee, S. Y. Kim, Choi, Chae

Critical revision of the manuscript for important intellectual content: all authors

Obtained funding: Chae, Choi

Study supervision: Chae, Choi

## Additional Contributions

We thank the patients, families and referring physicians who were involved in the study.

